# Detection, characterization, and phylogenetic analysis of a near-whole genome sequence of a novel astrovirus in an endemic Malagasy fruit bat, *Rousettus madagascariensis*

**DOI:** 10.1101/2023.10.27.564436

**Authors:** Sophia Horigan, Amy Kistler, Hafaliana Christian Ranaivoson, Angelo Andrianianina, Santino Andry, Gwenddolen Kettenburg, Vololoniaina Raharinosy, Tsiry Hasina Randriambolamanantsoa, Cristina M. Tato, Vincent Lacoste, Jean-Michel Heraud, Philippe Dussart, Cara E. Brook

## Abstract

Bats (order: *Chiroptera*) are known to host a diverse range of viruses, some of which present a public health risk. Thorough viral surveillance is therefore essential to predict and potentially mitigate zoonotic spillover. Astroviruses (family: *Astroviridae*) are an understudied group of viruses with a growing amount of indirect evidence for zoonotic transfer. Astroviruses have been detected in bats with significant prevalence and diversity, suggesting that bats may act as important astrovirus hosts. Most astrovirus surveillance in wild bat hosts has, to date, been restricted to single-gene PCR detection and concomitant Sanger sequencing; additionally, many bat species and many geographic regions have not yet been surveyed for astroviruses at all. Here, we use metagenomic Next Generation Sequencing (mNGS) to detect astroviruses in three species of Madagascar fruit bats, *Eidolon dupreanum, Pteropus rufus,* and *Rousettus madagascariensis*. We detect numerous partial sequences from all three species and one near-full length astrovirus sequence from *Rousettus madagascariensis*, which we use to characterize the evolutionary history of astroviruses both within bats and the broader mammalian clade, *Mamastrovirus*. Taken together, applications of mNGS implicate bats as important astrovirus hosts and demonstrate novel patterns of bat astrovirus evolutionary history, particularly in the Southwest Indian Ocean region.

## INTRODUCTION

Bats (order: *Chiroptera*) make up an extremely diverse mammalian order that has been identified as a reservoir of many of the world’s most virulent zoonotic viruses ^1–3^. Bats’ capacity to host virulent zoonotic viruses without experiencing disease is posited to be a byproduct of their evolution of flight, which necessitated metabolic adaptations leading to both elongated lifespans and immune system modifications promoting the evolution of viruses that are virulent to non-bat hosts^1,4–6^. Furthermore, the behavioral traits of many bat species, such as social grooming and roosting in dense aggregations, advance the transmission of infection within populations^7^. Despite the established importance of bats as viral reservoirs, existing surveillance of the viruses hosted by bats is uneven and shows marked geographic and taxonomic biases^8,9^.

A prime example is the island of Madagascar. It is home to 46 bat species with nearly 80% endemism, the result of a unique biogeographical history characterized by long isolation from surrounding landmasses^10,11^. This isolation has also led to the coevolution of extraordinarily diverse viruses within Malagasy bats^12–14^. Moreover, high human-bat contact rates due to hunting, particularly of large fruit bats, across the island increase the potential for zoonotic spillover^15^. Altogether, these factors make Madagascar an especially important location for surveillance of viruses harbored by bats.

A key understudied family of viruses are astroviruses (family: *Astroviridae*). Astroviruses (AstVs) are non-enveloped, positive-sense, single-stranded RNA viruses. Their genome contains a 5’-untranslated region (UTR), three open reading frames (ORFs)—ORF1a, ORF1b, and ORF2— and a 3’UTR with a poly A tail^16^. AstVs infect a remarkable diversity of hosts, grouping broadly into two genera: *Avastroviruses,* which infect avian hosts, and *Mamastroviruses,* which infect mammalian hosts, including humans. Approximately 2-9% of all acute non-bacterial gastrointestinal infections in children are thought to be from astrovirus infection^16,17^. This lack of host specificity demonstrates AstVs’ efficiency in cross-species transmission, thought to be facilitated by high viral diversity and frequent recombination events^16,18–21^. Although there is currently no direct evidence for zoonotic astrovirus transmission, a number of studies present indirect evidence suggesting a historical spillover of AstVs from animals to humans^20–25^10/27/23 2:23:00 PM. Despite their clear importance in both wildlife and public health, we have only recently begun to understand the true diversity of astroviruses and their evolutionary history in non-human hosts.

Astroviruses have been detected at high incidence and with remarkable diversity in both suborders of *Chiroptera*, Yinpterochiroptera and Yangochiroptera, though these values range widely depending on the study design, species, and location^24–28^. Differing phylogenetic analyses reveal both strong and weak host- and geographic-clustering, with some bat AstVs grouping more closely with members of the avian *Avastrovirus* genus. High astrovirus diversity detected within members of the same populations of bat hosts suggests the simultaneous circulation of multiple strains, perhaps driven by the co-roosting of multiple bat species in dense aggregations^1,7,29^. Altogether this evidence suggests that bats may be an important reservoir source for cross-species transmission or zoonotic spillover of astroviruses. These insights, however, have been largely based on PCR detections and single gene sequences (typically the RNA-dependent RNA polymerase gene) derived from a few species of bats, limiting the potential for more robust evolutionary analysis. Very recently, metagenomic next-generation (mNGS) sequencing has resulted in the detection of five full-genome bat astroviruses globally^30–32^. However, many outstanding questions remain regarding astrovirus presence, infection dynamics, and zoonotic risk.

Here, we present mNGS detection and characterization of astroviruses sampled from three species of endemic fruit bat from Madagascar*: Pteropus rufus, Rousettus madagascariensis,* and *Eidolon dupreanum*. We identified a novel near-full length astrovirus genome detected in a *Rousettus madagascariensis* fecal sample and we characterized its evolutionary history among the broader clade of *Mamastroviruses.* Using the RNA-dependent RNA polymerase (RdRp) conserved region of this genome, we additionally explored the biogeographical history of bat astroviruses from Madagascar and its surrounding landmasses in the Southwest Indian Ocean Region, shedding light on the evolutionary history of astroviruses between bat suborders, Yinpterochiroptera and Yangochiroptera.

## METHODS

### Sample collection and processing

Astrovirus infections were identified from a dataset of viruses detected in samples from a longitudinal study of fruit bats across Madagascar. Methodological details on bat field sampling and subsequent RNA extraction, library preparation, and Illumina sequencing have been reported in previous work;^13,14^ here, we give only a brief overview.

Between 2018 – 2019, monthly bat captures were carried out at four species-specific locations: Ambakoana roost (−18.513 S, 48.167 E, *Pteropus rufus)*; Angavobe cave (−18.944S, 47.949 E, *Eidolon dupreanum)*; Angavokely cave (−18.933 S, 47.758 E, *Eidolon dupreanum)*; Maromizaha cave (−18.9623 S, 48.4525 E, *Rousettus madagascariensis).* Bats were identified to species, sex, and age (adult vs juvenile), and throat, fecal, and urine samples were collected.

Following field collection, throat, fecal, and urine samples underwent RNA extraction in the Virology Unit at the Institut Pasteur de Madagascar (IPM) using the Zymo Quick DNA/RNA Microprep Plus Kit (Zymo Research, Irvine, CA). In total, RNA from 285 fecal, 143 throat, and 196 urine swab samples was extracted, then stored in −80^0^ freezers at IPM, prior to final transport on dry ice to the Chan Zuckerberg Biohub San Francisco (CZB-SF) for eventual library preparation and subsequent mNGS.

Aliquots of each sample were arrayed into a 384 well plate for mNGS library preparation. Samples were evaporated using a GeneVac EV-2 (SP Industries, Warminster, PA, USA) to enable miniaturized library preparation with the NEBNext Ultra II RNA Library Prep Kit (New England Biolabs, Beverly, MA, USA). Library preparation was performed per the manufacturer’s instructions, with the following modifications: 25 pg of External RNA Controls Consortium Spike-in mix (ERCCS, Thermo-Fisher) was added to each sample prior to RNA fragmentation; the input RNA mixture was fragmented for 8 min at 94^0^C prior to reverse transcription; and a total of 14 cycles of PCR with dual-indexed TruSeq adaptors was applied to amplify the resulting individual libraries. Samples were assessed for quality and quantity, then submitted to an Illumina NovaSeq (Illumina, San Diego, CA, USA) for paired-end sequencing (2 x 146 bp). The pipeline used to separate the sequencing output of the individual libraries into FASTQ files of 146bp paired-end reads is available on GitHub at https://github.com/czbiohub/utilities.

### Detection

Raw reads from Illumina were host-filtered, quality-filtered, and assembled on the Chan Zuckerberg Infectious Diseases (CZID) bioinformatics platform (v3.10 NR/NT 2019-12-01)^33^, using a host background model of “bat” compiled from all publicly available full-length bat genomes in GenBank at the time of sequencing (July 2019). Samples were marked positive for astrovirus infection if at least two contigs with an average read depth >2 reads/nucleotide were assembled that showed significant nucleotide or protein BLAST alignment(s) (alignment length >100nt/aa and E-value < 0.00001 for nucleotide BLAST/bit score >100 for protein BLAST) to astroviruses present in NCBI NR/NT database (v12-01-2019). Additionally, all non-host contigs assembled in CZID were manually BLASTed against all full-length and protein reference sequences for astroviruses available in NCBI Virus.

To test for differences in astrovirus prevalence, we performed four Pearson’s Chi squared tests between differing subsets of the data: between total prevalence across the three species, and between adults and juveniles within each species.

### Genome Annotation and % Identity Analysis

To annotate coding sequences, we downloaded all available bat astrovirus full genomes from NCBI Virus at the time of analysis (August 2022). We aligned positive astrovirus contigs from our dataset to these background sequences using the MAFFT^34^ algorithm (v7.450) with default parameters in Geneious Prime (v08-18-2022). We then annotated open reading frames and genes in our novel sequence by identifying stop and start codons in regions adjacent to those identified in the homologs.

To investigate similarity to other published sequences, we performed BLASTn (nucleotide-nucleotide) and BLASTx (translated nucleotide-protein) searches within the NCBI database (Table S1). Additionally, we created amino acid and nucleotide identity plots using the program pySimplot^35^ with input alignment generated using MAFFT^34^ (v7.450) with default parameters in Geneious Prime (v08-18-2022).

We identified one near-full length astrovirus genome within this dataset which was used for all subsequent phylogenetic analyses. All other astrovirus hits were short fragments which aligned with regions with limited phylogenetic potential given the lack of PCR targeting of that region in other studies.

### Phylogenetic Analysis

To perform phylogenetic analysis, we combined our novel near-full length sequence with those publicly available on NCBI. We carried out three major phylogenetic analyses, building (a) a full-genome *Mamastrovirus* maximum likelihood (ML) phylogeny, (b) a time-resolved Bayesian phylogeny corresponding to a selection of full genome *Mamastrovirus* sequences available on NCBI Virus, and (c) a *Mamastrovirus* ML phylogeny corresponding to a conserved 410 bp fragment of the RNA-dependent RNA polymerase (RdRp) gene encapsulated in the AstV ORF1b with a focus in the South West Indian Ocean region. Detailed methods for the construction of each phylogeny are available on GitHub (see Data Availability).

### Sequence Compilation

Our full genome ML phylogeny consisted of one novel full length *Mamastrovirus* sequence from our study, combined with 41 unique *Mamastrovirus* sequences from NCBI, and one full length *Avastrovirus* sequence as an outgroup, for a total of 43 sequences. We compiled the *Mamastrovirus* sequences from NCBI through three queries, selecting: [A] all complete RefSeq Genomes under Virus: *Mamastrovirus* (taxid:249588) and Virus: *unclassified Mamastrovirus* (taxid:526119) greater than 6,000 bp (N=36), [B] *Mamastrovirus* nucleotide genomes under Virus: *Astroviridae* (taxid:39733) and Virus: *unclassified Astroviridae* (taxid:352926) with Host: *Chiroptera (bats)* (taxid:9397) over 6,000 bp (N=2), and [C] manual searching of *Mamastrovirus* nucleotide genomes >6,000 bp identified in the literature (N=3).

Our Bayesian timetree consisted of the same set of full length *Mamastrovirus* sequences, removing the *Avastrovirus* outgroup, for a total of 42 sequences.

Our *Mamastrovirus* RdRp ML phylogeny consisted of an overlapping 410 bp fragment in the center of the RdRp region of our one full length *Mamastrovirus* sequence, combined with 122 unique bat *Mamastrovirus* sequences from NCBI, and one Avastrovirus RdRp fragment as an outgroup, for a total of 124 sequences. NCBI sequences were restricted to those from bat hosts sampled in the Southwest Indian Ocean (SWIO) region. They were compiled through one query in NCBI Virus: Virus: Astroviridae (taxid:39733), Host: *Chiroptera* (taxid:9397), and Geographic Region: Madagascar (64), Mozambique (31), and Reunion (27). No sequences were available from the Comoros, Mayotte, Mauritius, and the Seychelles. Sequences were confirmed to be RdRp fragments via alignment, and metadata such as host taxa and sampling location were verified in the source literature.

### Alignment and Substitution Model

Following dataset compilation for each phylogenetic analysis, sequences were aligned via the MAFFT^34^ (v7.450) algorithm in Geneious Prime (v 2022-08-18) using default parameters. Alignments were visually examined and trimmed to match the shortest sequence in the dataset. We then used Modeltest-NG^36^ (v0.1.7) to determine the best fit nucleotide substitution for each alignment. All sequences, subsets, and alignments are available in our open-source GitHub repository (see Data Availability).

### Phylogenetic Tree Assembly

Both the full genome and RdRp ML trees were constructed in RAxML-NG^37^ (v1.1.0), using the best nucleotide substitution model from Modeltest-NG^36^. Following best practice recommendations in RAxML-NG^37^, twenty ML inferences were made, followed by bootstrap replicate trees inferred using Felsenstein’s method^38^. The MRE-based bootstrapping test was performed every 50 replicates, and bootstrapping was terminated when the diagnostic result was below the threshold value. Support values were compiled onto the best-scoring tree.

The Bayesian timetree was built in BEAST2^39^ (v2.6.7), using the best nucleotide substitution model from Modeltest-NG^36^. We used a Bayesian Skyline Coalescent Model with a strict lognormal clock rate with prior mean 0.001 substitutions/site/year^40^, and a constant population size. Sampling date for each sequence was inferred from NCBI ‘Collection Date’ or though reading source literature; if day was not available, the sampling date was set to the 15^th^ of the month listed; if day and month were not available, the sampling date was set to July 15^th^ of the collection year. Markov Chain Monte Carlo (MCMC) chains were run for >700,000,000 iterations and terminated when we identified convergence at ESS values > 200 using TRACER (v1.7), with 10% burn-in. We used TreeAnnotator (v2.6.3) to examine mean posterior densities at each node.

All phylogenies were visualized in RStudio (v2022.07.01), using the package ‘ggtree’^41^.

### Nucleotide Sequence Accession Numbers

One annotated near full-length genome sequence from *Rousettus madagascariensis* was submitted to NCBI and is available under accession number OQ606244.

## RESULTS

### Detection

In total, RNA samples from 285 fecal, 196 urine, and 143 throat swabs were sequenced. Astrovirus positives were defined as samples harboring two or more contigs with an average read depth >2 reads with statistically significant blastn or blastx alignment to astrovirus sequences in NCBI (alignment length >100 nt/aa and E-value < 0.00001 for nucleotide BLAST/bit score >100 for protein BLAST). Based on these criteria, evidence of astrovirus infection was detectable in all 3 species of Malagasy bats, across all sampled locations (Figure 1). Astrovirus positives were detected among 4/285 (1.4%) fecal samples and 7/196 urine samples (3.6%), and none of the 143 throat swab samples.

**Figure 1:**
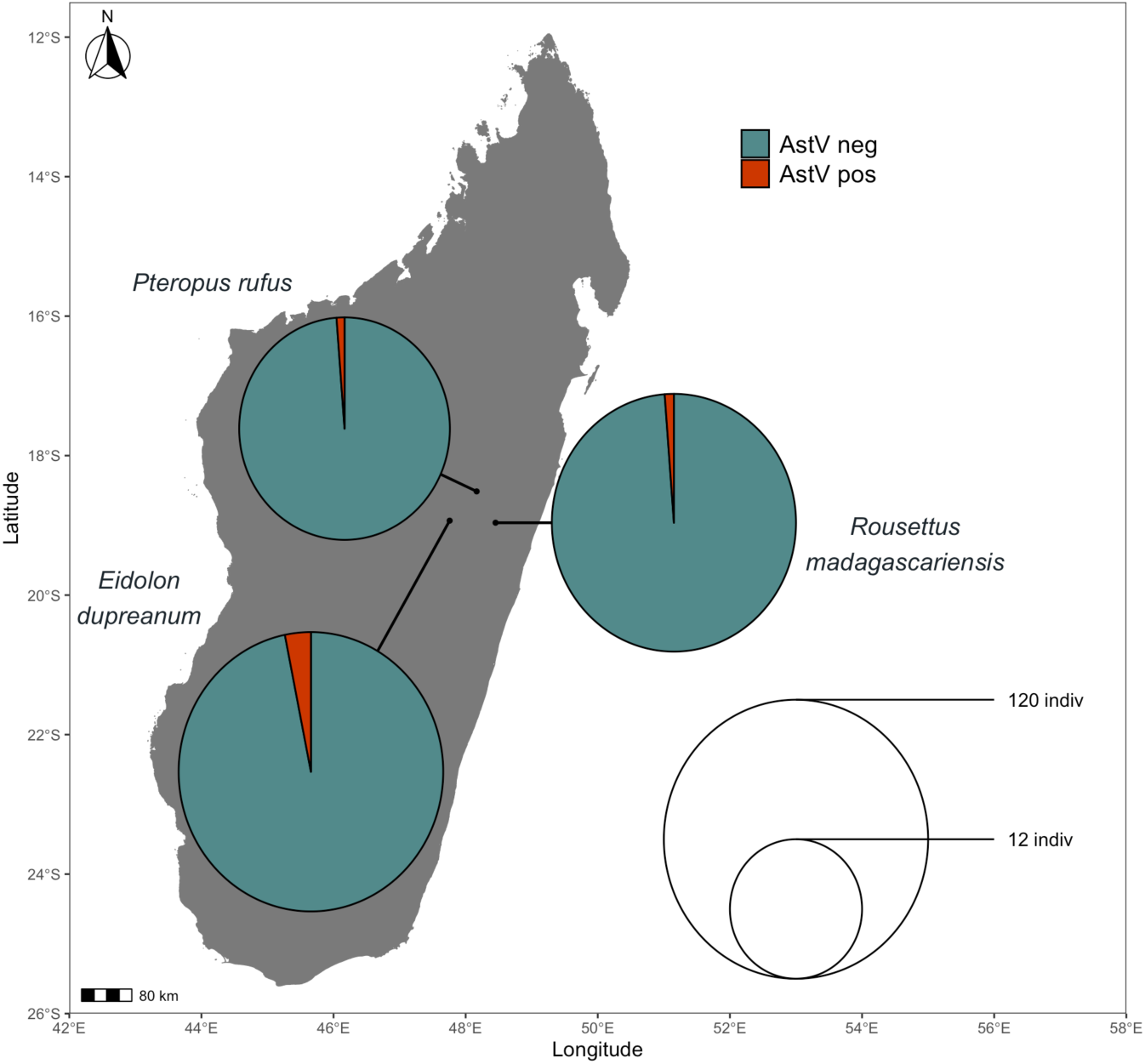
Map of sampling sites for *P. rufus*, *E. dupreanum*, and *R. madagascariensis* in the districts of Moramanga and Manjakandriana, Madagascar (*P. rufus:* Ambakoana roost; *E. dupreanum*: Angavobe/Angavokely caves; *R. madagascariensis*: Maromizaha cave). Pie charts correspond to astrovirus prevalence of any sample type in bats across all three species: 1/44 (2.27%) for *P. rufus*, 8/145 (5.52%) for *E. dupreanum*, and 2/96 (2.08%) for *R. madagascariensis*. Pie circle size corresponds to sample size on a log-10 scale.

Prevalence varied slightly between species, with 1/44 (2.3%) *Pteropus rufus*, 8/145 (5.5%) *Eidolon dupreanum*, and 2/96 (2.1%) *Rousettus madagascariensis* individuals identified for positive astrovirus infection in either fecal or urine samples (Figure 1). These between-species differences were not significant (χ^2^=2.104, P=0.3492). Juvenile vs adult prevalence also did not vary significantly: 1/15 (6.7%) vs. 0/29 (0%) for *P. rufus* (χ^2^=1.941, P=0.1635), 1/13 (7.7%) vs. 7/132 (5.3%) for *E. dupreanum* (χ^2^=0.122, P=0.7264), and 1/13 (7.7%) vs. 1/83 (1.2%) for *R. madagascariensis* (χ^2^=2.271, P=0.1318).

### Sequence Analysis of Malagasy AstVs

In total, 21 astrovirus contigs were identified, ranging from 209 – 6,593 base pairs in length. The longest contig, AstV OQ606244, recovered from a fecal sample from a juvenile *Rousettus madagascariensis*, had substantial supporting read depth (Figure 2), encoding a near complete genome sequence. Basic open reading frame analysis and sequence alignment to *Mamastroviruses* genomes from NCBI revealed that the entirety of ORF1b and ORF2 are present in the near full genome *R. madagascariensis* AstV. ORF1a is nearly fully represented in the contig; however, a fragment of the 5’ region including the start codon was not captured.

**Figure 2:**
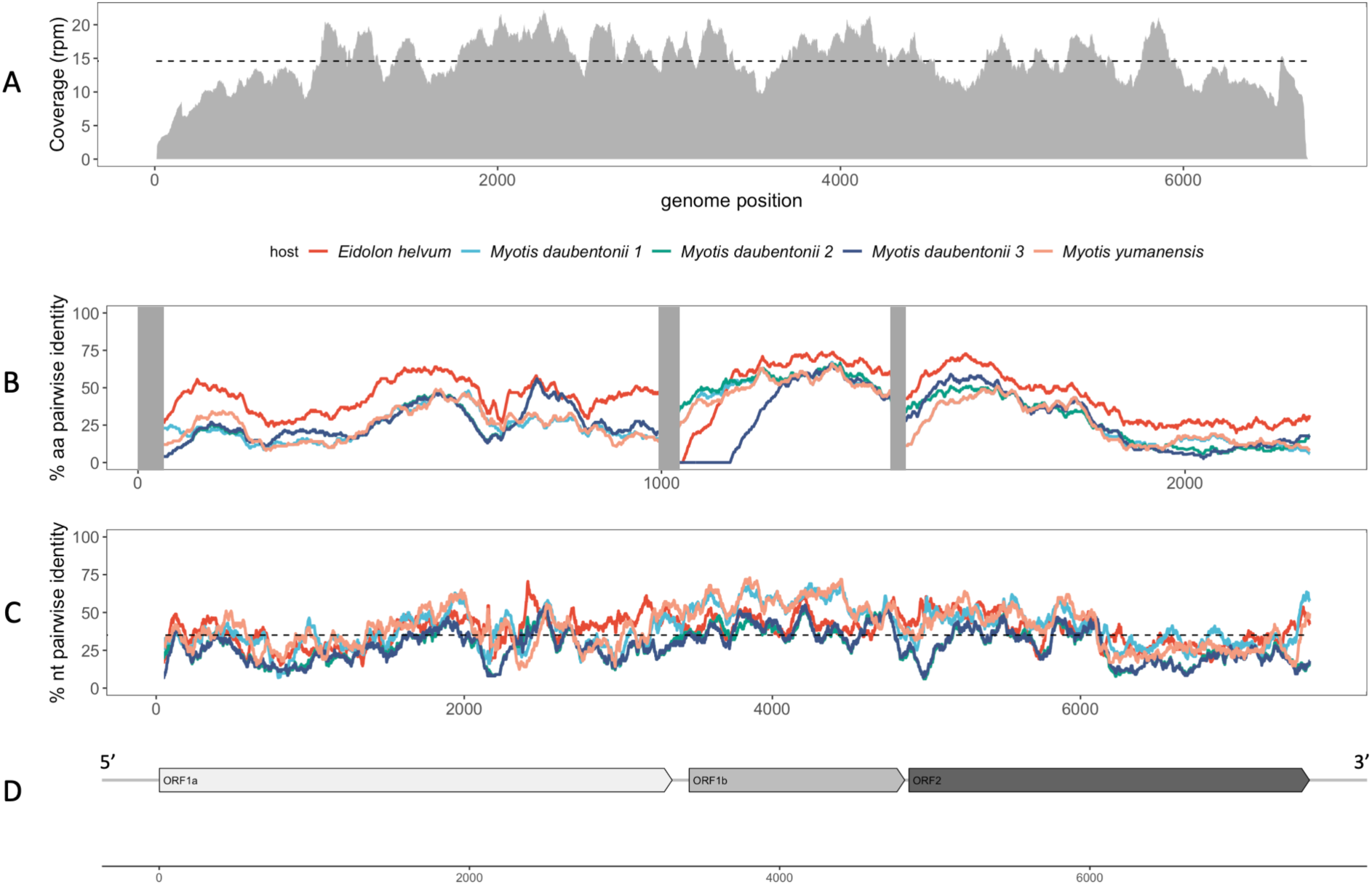
Genomic structure of Astroviruses. Identity plots comparing AstV OQ606244 sequence with all available chiropteran full genome astrovirus sequences on NCBI. Top panel depicts amino acid % identity, and middle panel depicts % nucleotide identity. In both panels, the query sequence is *R. madagascariensis* AstV (OQ606244), and comparison sequences are: MG693176 *(Eidolon helvum)*, MZ218054 *(Myotis daubentoniid 1),* MZ218053 *(Myotis daubentoniid 2),* MN832787 *(Myotis daubentoniid 3),* MT734809 *(Myotis yumanensis).* Bottom panel depicts a coverage plot showing contig depth (reads per million from total reads recovered for this sample) for assembled full-genome OQ606244.

All other astrovirus detections from our sampling produced contigs representing fragments that aligned within the ORF2 region. Because this region is not typically targeted in PCR, there were not enough NCBI submissions available for a thorough evolutionary analysis including these fragments. As a result, these positive detections were not analyzed further. BLAST analysis of the near-full AstV genome contig (NCBI Accession # OQ606244) demonstrated that it is highly divergent from previously described Mamastrovirus sequences. BLASTn query of AstV OQ606244 recovered identity with only one other sequence in NCBI: a small portion (5%) of AstV OQ606244 could be aligned to a Mamastrovirus RNA-dependent RNA-polymerase (RdRp) from an unknown Chiropteran species sampled in Tanzania (KY054020) at 80.9% identity. BLASTx analysis of the highest scoring matches in the database for the amino acid sequence of AstV OQ606244 revealed that the best amino acid alignment shared only 43.1% amino acid identity across 40% of an AstV ORF1a protein sequence recovered from *Eidolon helvum* in Cameroon (MG693176). All other top results were to non-chiropteran species including pigs and raccoon dogs (Table S2).

To further investigate the sequence divergence of AstV OQ606244, we performed scanning pairwise nucleotide and amino acid sequence analysis using AstV OQ606244 paired, in turn, with each of five full genome bat Astrovirus sequences available in NCBI (Figure 2). We observed low nucleotide sequence identity: from 28.18% - 40.28% across the whole genome, and average amino acid identity from 23.87% and 44.29% in ORF 1a, 35.14% - 55.61% in ORF1b, and 24.84% - 40.73% in ORF2 (Table 1).

**Table 1:** Summary table of Figure 2 scanning pairwise identity plots showing % identity of each queried astrovirus to AstV OQ606244 across whole-genome nucleotide sequence and amino acid open-reading frames (ORF) 1a, 1b, and 2.

| NCBI ID | Host Species | % nucleotide identity | % ORF1a aa identity | % ORF1b aa identity | % ORF2 aa identity |
| --- | --- | --- | --- | --- | --- |
| MG693176 | *Eidolon helvum* | 38.84 | 44.29 | 55.61 | 40.73 |
| MN832787 | *Myotis daubentonii* | 39.84 | 23.87 | 51.78 | 24.84 |
| MT734809 | *Myotis daubentonii* | 40.28 | 24.78 | 50.31 | 24.92 |
| MZ218053 | *Myotis daubentonii* | 28.18 | 25.99 | 54.26 | 25.59 |
| MZ218054 | *Myotis yumanensis* | 28.24 | 25.92 | 35.14 | 26.65 |

We observed slight discordance in the best matches with AstV OQ606244 at the nucleotide and amino acid level in this pairwise sequence analysis with complete AstV genomes identified in other chiropteran species. One astrovirus (MT734809) from one of the four *Myotis daubentonii* hosts had the highest pairwise average nucleotide identity to AstV OQ606244. In contrast, a different AstV genome identified in *Eidolon helvum* (MG693176)—which, like *R. madagascariensis*, is a Pteropodid fruit bat—had the highest pairwise amino acid identity to AstV OQ606244 across all open reading frames.

### Phylogenetic Analysis

### *Mamastrovirus* full-genome evolutionary history

To investigate the evolutionary history of our novel bat astrovirus within the broader genus of mammalian astroviruses, *Mamastrovirus,* we built a maximum-likelihood phylogenetic tree including RefSeq genomes from a range of hosts with an avian astrovirus from the genus *Avastrovirus* as an outgroup. The best fit nucleotide substitution model as generated by Modeltest-NG^36^ was TVM+I+G4. The full-genome maximum likelihood tree resolved two distinct *Mamastrovirus* clades, with a single Porcine astrovirus^42^ falling out more closely related to the *Avastrovirus* outgroup, likely because it falls within a clade not represented in this phylogeny (Figure 3A). No astroviruses derived from any single host order demonstrated monophyly, though sequences nonetheless clustered based on host taxonomy, with high support in most cases. Overall, these findings lend additional support to a growing body of evidence suggesting that astroviruses are likely to engage in cross-species transmission^16,18–21^.

**Figure 3:**
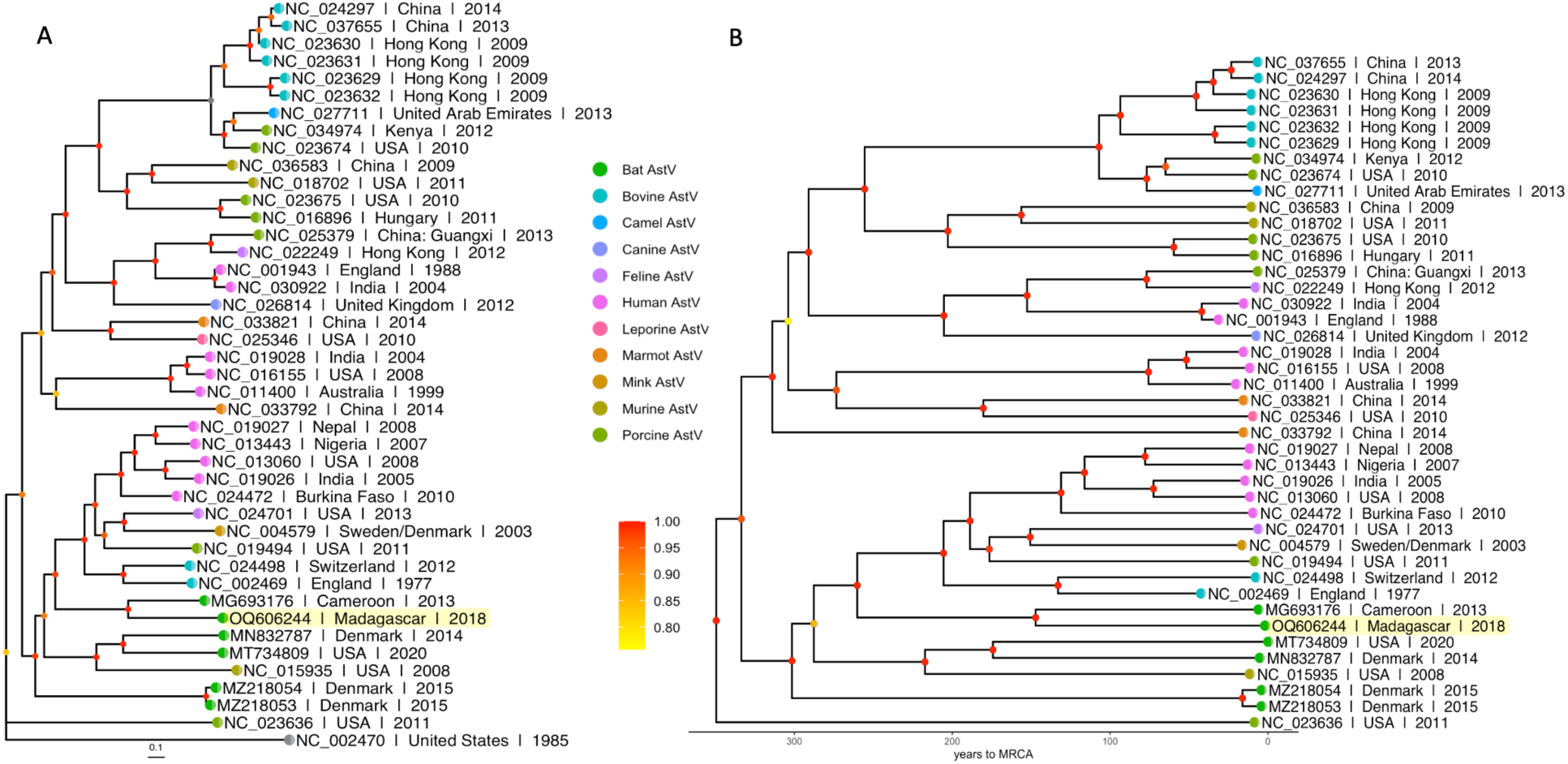
**A)** Maximum likelihood phylogeny of full genome *Mamastrovirus* sequences (RAxML-NG, TVM+I+G4). Bootstrap values computed using Felsenstein’s method^38^ are visualized on tree branches. Tree is rooted by a turkey *Avastrovirus* (NC_002470) and a divergent Porcine *Mamastrovirus* (NC_023636). Branch lengths are scaled by nucleotide substitutions per site, corresponding to the scalebar. **B)** Bayesian time-resolved phylogeny of full genome *Mamastrovirus* sequences generated from >700,000,000 steps under a Bayesian Skyline Coalescent Model (TVM+I+G4). Node color represents mean posterior estimates averaging over all steps with 10% burn-in (see scale bar on left). **A,B)** Tip labels include NCBI taxon ID, strain, host species, location of collection, and year of collection. Tip points are colored by astrovirus strain, with AstV OQ606244 highlighted in yellow.

The bat astroviruses formed a paraphyletic group, with two sequences basal to a broad clade containing a collection of murine, bovine, porcine, feline, and human astroviruses. AstV OQ606244, sampled from *R. madagascariensis*, grouped most closely with a previously described sequence recovered from a Cameroonian *Eidolon helvum* fruit bat, a fellow Yinpterochiropteran host^30^. In our phylogeny, both *R. madagascariensis* and *E. helvum* astroviruses resolved as more recently derived than four previously described full genome astroviruses recovered from Yangochiropteran (*Myotis* spp.) hosts (Figure 3A).

We recovered largely the same topology in our time-resolved Bayesian phylogeny as in the ML tree: all sequences grouped into same two main *Mamastrovirus* clades, with a few slight deviations in the sequences that resolved as basal within each subclade (Figure 3B). All bat astroviruses resolved the exact same position in both the Bayesian and ML tree, reinforcing resolution of their evolutionary positions. Our Bayesian timetree recovered a relatively shallow evolutionary history for the *Mamastrovirus* clade, dating the most recent common ancestor (MRCA) of all Mamastroviruses to just over 300 years ago, and the MRCA of our *R. madagascariensis* astrovirus and its closest *E. helvum* AstV relative ∼150 years ago.

To investigate if bat astrovirus evolutionary history is driven by biogeographical patterns or host relationships, we built a maximum likelihood phylogeny of all known bat astrovirus RdRp sequences from the Southwest Indian Ocean region. Focusing on the RdRp region greatly expanded the range of background sequences available for inclusion in phylogenetic analysis, as this region of the AstV genome is the most commonly targeted by single-gene PCR detection methods. In this phylogeny, the best fit nucleotide substitution model as generated by Modeltest-NG^36^ was TVM+I+G4. Astrovirus sequences in the RdRp phylogeny grouped largely according to host taxonomy rather than geography of sampling site, a pattern which held across both suborder and family across all sampling locations. Almost all Yinpterochiropteran *Mamastrovirus* sequences resolved into a monophyletic group, while Yangochiropteran AstVs grouped into multiple clades, all basal or sister to the single Yinpterochiropteran clade.

Within Yangochiropteran hosts, AstV sequences derived from Molossids and Miniopterids clustered together regardless of location. Molossid-derived AstVs resolved into numerous scattered clades basal to all other known sequences. Paraphyly has been detected in Molossid species genomes^43^, and thus may promote paraphyly in their viruses through coevolution. This host restriction, however, may be confounded with sampling location, since only Reunion Island Molossids and Madagascar Miniopterids were represented in this analysis. Notably, previous sampling from Madagascar Molossids has, to date, detected no astroviruses^12^ (Figure 4); however this previous sampling did not include all Molossid populations or species across the island.

**Figure 4:**
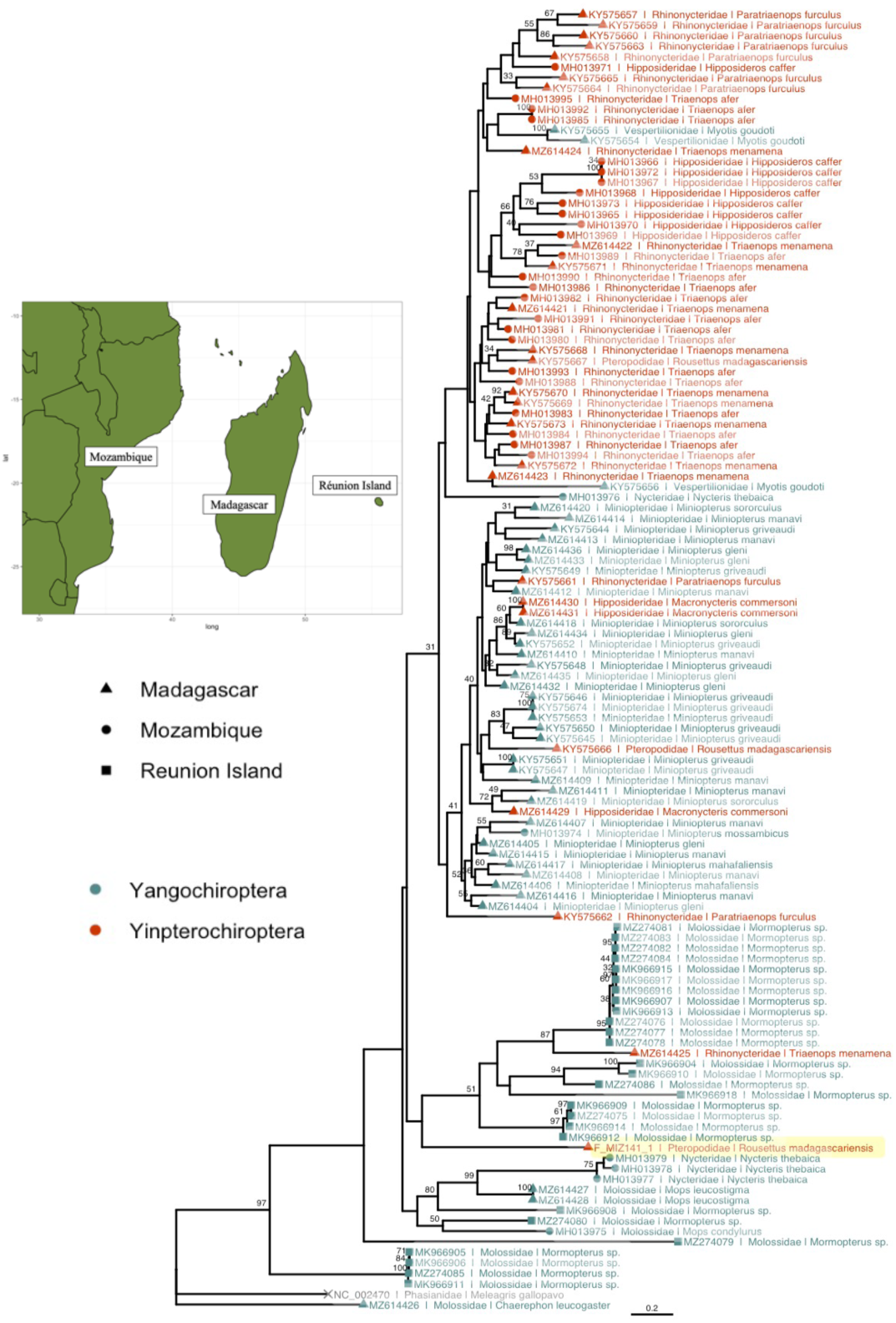
**A)** Map showing locations in the Southwest Indian Ocean (SWIO) region with bat astrovirus samples. Map generated using the ‘maps’ package in R Studio. **B)** Maximum Likelihood phylogeny of a 401bp fragment of *Mamastrovirus* Orf1b (RdRp) from Southwest Indian Ocean (SWIO) bat hosts (RAxML-NG, TVM+I+G4). Bootstrap values computed using Felsenstein’s method^38^; values >30 are visualized on tree branches. Tip labels include NCBI taxon ID, host family, and host species. Tip points are colored by sub-order of bat host and shaped by country of origin. AstV OQ606244 is highlighted in yellow. Tree is rooted by a Turkey *Avastrovirus* (NCBI taxID: NC_002470) and a divergent bat *Mamastrovirus* (NCBI taxID: MZ614426). Branch lengths are scaled by nucleotide substitutions per site, as indicated by the scale bar.

There were a few notable exceptions to this pattern. AstVs recovered from Nycterid (family: Nycteridae) bats resolved into two disparate locations across the phylogeny: one clustered within the Molossids and another as basal to the Yinpterochiropteran clade. Vespertilionid AstVs (family: Vespertilionidae), with a sample size of only two, were placed within the Yinpterochiropteran clade, most closely related to Rhinonycterid AstVs (family: Rhinocyteridae).

Within the Yinpterochiropteran AstVs, Rhinonycterid astroviruses mixed with those recovered from Hipposiderids, both with representatives from Madagascar and Mozambique. Numerous prior publications have established these bat families as sister to one another^43,44^; the genetic similarity of these host clades makes cross-transmission of viruses between them more likely.

As the only AstV representatives in this phylogeny hosted by bats in the Pteropodidae family, three AstV sequences from *Rousettus madagascariensis* did not group together. All three resolved to very different places on the phylogeny—one within the Yinpterochiropteran clade and most closely related to a Rhinonycterid AstV, one within the Yangochiropteran Miniopterid clade, and AstV OQ606244 within the Yangochiropteran Molossid clade.

Altogether, these patterns suggest that the Yinpterochiropteran AstV sequences that resolve within the Yangochiropteran AstV clade could represent cross-species spillovers. The fact that there are more of these putative spillover AstV sequences from Yangochiropteran to Yinpterochiropteran hosts than the reverse in our phylogeny—as well as the basal position of the Yangochiropteran relative to Yinpterochiropteran AstV clade—suggests that SWIO bat AstVs may have originated Yangochiropteran hosts, then spilled over and radiated within Yinpterochiropteran hosts. Nonetheless, the weak node support and lack of representation of many bat species in this tree preclude certainty in this conclusion.

## DISCUSSION

### Detection, Divergence, and Zoonotic Potential

Here we describe how mNGS sequencing was applied to detect and characterize astroviruses from three endemic species of Malagasy fruit bat, *Pteropus rufus, Rousettus madagascariensis,* and *Eidolon dupreanum.* We characterize the first near-full genome astrovirus detected from *Rousettus madagascariensis* and undertake phylogenetic analysis of the surrounding clade.

In addition, this work presents the first detection of astroviruses from *Eidolon dupreanum* and *Pteropus rufus*. Astroviruses have been detected previously in *Rousettus madagascariensis*^12^, though we contribute the first known near-full length genome sequence of this virus family from this host. Astrovirus RNA has now been identified in 14/18 species of Malagasy bats that have been queried for this virus family (Table S3). While we did detect astrovirus infection in representatives from each of our three sampled species, our detected prevalence was (< 8%). It is possible that this low rate of astrovirus detection could reflect a reduced sensitivity of mNGS for virus detection; however, similarly low AstV prevalence levels have also been described in other bats, particularly Pteropodids^45,46^ when more sensitive directed detection methods were used. Previous PCR-based sampling of astroviruses in *Rousettus madagascariensis* found positive infection in 2/41 (4.8%)^12^ and 2/40 (5.0%)^47^, comparable to our 2.1%. Detection of astroviruses in other species of Malagasy bats, however, has reached as high as 88.9%^47^ prevalence, but these high values appear to be more represented in Yangochiropteran bats.

Both open-ended BLAST and our directed, pairwise sequence comparison of one near full-length *R. madagascariensis* AstV genome with all other bat astrovirus genomes available indicate that this newly detected AstV is significantly diverged from all previously described viruses in the family (Table S1, Figure 2). Previous genomic analyses using this same mNGS dataset have also demonstrated the extraordinary divergence of multiple other Madagascar bat viruses^13,14^. Kettenburg et al.^13^ (2022) identified a novel *Nobecovirus* (family: *Coronaviridae*) in a *Pteropus rufus* fecal specimen which is highly divergent from all others known and appears to be basal to the entire *Nobecovirus* subgenus of the Betacoronavirus clade^13^. Madera et al.^14^ (2022) described a novel *Henipavirus* (family: *Paramyxoviridae)* from *Eidolon dupreanum* urine samples, which is highly diverged from all other viruses in its genus and also basal to those known to be hosted by bats. It is likely that future sampling of the Malagasy bat virome will continue to reveal more highly divergent viruses, reflecting Madagascar’s long geographic isolation from other land masses.

Intriguingly, AstV OQ606244, demonstrates divergence from previously characterized bat astroviruses in its ORF2 region (Figure 2). Because ORF2 encodes the AstV spike protein, which mediates virus entry into host cells, this region is vital to understanding potential cross-species transmission^16^. While the precise mechanisms of astrovirus cell entry are unknown, the divergence of AstV OQ606244 from other astroviruses in this region highlights the need for *in vitro* studies to quantify zoonotic potential in this virus family. Altogether, the highly divergent sequence of AstV OQ606244, paired with high observed human-bat contact rates in Madagascar, underline the need for heightened astrovirus surveillance in this region.

### Evolutionary History

Our full-genome ML and Bayesian phylogenies shed light on the placement of bat astroviruses with respect to other mammalian astroviruses. In our phylogenetic analyses, two previously described bat astrovirus genome sequences resolved as basal sequences within one of the two distinct *Mamastrovirus* clades. This basal position indicates that bats may have been important ancestral hosts from which other mammalian astroviruses descended. We estimate a relatively shallow evolutionary history for the entire *Mamastrovirus* clade, with MRCA dating to only ∼300 years ago. Given the substantial host diversity of astroviruses, this indicates that astroviruses circulate widely within mammalian populations. Our phylogenetic trees also give us insight into astrovirus evolutionary history within bats. Our tree topologies suggest that astroviruses may have originated in insectivorous Yangochiropteran bats and later been transmitted to and further radiated in Yinpteropchiropterans.

We date the MRCA of AstV OQ606244 and its most closely related known relative, an *Eidolon helvum* AstV, to ∼150 years ago. Considering Madagascar’s isolation from the African continent for ∼160 million years and the divergence of the *Rousettus* spp. genus ∼20 million years ago^48^, this would indicate very recent viral genetic exchange between bats from Madagascar and the African continent. Thirty-eight of Madagascar’s 46 bat species are endemic, while two species exhibit ranges that include the Comoros and Reunion Islands (*Scotophilus borbonicus* and *Miniopterus aellini*, respectively), and nine species can be found more broadly distributed across Africa, and, in some cases, parts of Asia and Europe (e.g. *Pipistrellus kuhlii* and *Mops midas)*^10^. Investigating which astroviruses are detectable in these cosmopolitan species presents a key opportunity to elucidate the role of these more broadly distributed potential host species in the diversification and distribution of astroviruses.

In contrast to many other astrovirus studies, which demonstrate minimal to no host restriction on AstV evolution^12,46,49–52^, our RdRp phylogeny, which contained representative AstV sequences from SWIO region bats, indicated that astroviruses group by host phylogeny rather than by sampling location^43^. Perhaps the unique biogeographic features of the SWIO region, whereby island status limits bat dispersal, makes this pattern more likely. The most notable exception to this pattern is the appearance of eight Yinpterochiropteran sequences interspersed within disparate Yangochiropteran clades, in contrast with only three Yangochiropteran sequences clustered within the single Yinpterochiropteran clade. For bats, cross-species virus transmission can be mediated by interspecies co-roosting ^1,4,7,53^. Both *Rousettus madagascariensis* and *Eidolon dupreanum*, cave-roosting species, have been noted to co-roost with several species of insectivorous bats, which may facilitate cross-species transmission^10^. Indeed, Molossid species bats (*Mormopterus jugularis)* have been described co-roosting with *R. madagascariensis* in the exact same roost from which our AstV sequence was recovered^53^, and our *R. madagascariensis* astrovirus clusters within a clade otherwise characterized by Molossid-derived AstVs.

It should be noted, however, that the relatively short fragment length of the RdRp region in our phylogenetic analysis (approximately 401 bases) yields weak support across many nodes, particularly those that are basal. Furthermore, sampling has been historically biased to favor insectivorous bats in the SWIO region; as a result, the lack of representation of many bat host species in AstV surveillance to date limits our understanding. Given the high divergence of our recovered AstV sequence, however, as well as the propensity for human-bat interactions in the SWIO region, further surveillance of previously unsampled species to fill these astrovirus research gaps represents a major public health priority.

## Conclusion

We characterize astroviruses recovered from urine and fecal samples derived from three species of Malagasy fruit bats, *Pteropus rufus*, *Eidolon dupreanum*, and *Rousettus madagascariensis*, identifying one near-full length astrovirus genome from *Rousettus madagascariensis*. This virus is highly divergent from all previously described bat AstV sequences, reflecting Madagascar’s unique biogeographical history as an island nation. This work supports the growing body of literature demonstrating that bats are likely to host considerable astrovirus diversity. Given the high rates of bat-human contact in this region this work demonstrates the urgency in the surveillance of astroviruses and other viruses of zoonotic potential in Malagasy bats, particularly. We advocate particularly for the role of longitudinal studies in addressing these aims, given the potential to develop a nuanced understanding of temporal and spatial dynamics in viral presence and shedding. We additionally advocate for heightened characterization of more whole genome bat astrovirus sequences to strengthen downstream phylogenetic analyses.

## ACKNOWLEDGEMENTS

The authors acknowledge Anecia Gentles, Kimberly Rivera, Fifi Ravelomanantsoa, and Sarah Guth for help in the field and lab. We acknowledge the Virology Unit at the Institut Pasteur de Madagascar and the Department of Zoologie et Biodiversité Animale at the University of Antananarivo. We thank Joseph L. DeRisi, Maira Phelps, and Vida Ahyong of the Chan Zuckerberg Biohub (CZB) for logistical support. We additionally thank Angela Detweiler, Michelle Tan, and Norma Neff of the CZB genomics platform for mNGS support and thank the Brook lab at the University of Chicago for helpful contributions to the manuscript. This work was completed in part with resources provided by the University of Chicago’s Research Computing Center.

## DATA AVAILABLITY STATEMENT

The near-full length genome presented in the study is deposited in NCBI, accession number: OQ606244. Detailed methods are available at https://github.com/brooklabteam/Mada-Bat-AstV.

## ETHICS STATEMENT

The animal study was reviewed and approved by UC Berkeley Animal Care and Use Committee and Madagascar Ministry of Forest and the Environment under guidelines posted by the American Veterinary Medical Association.

## AUTHOR CONTRIBUTIONS

CEB conceived of the project and acquired the funding, in collaboration with J-MH, VL, and PD. Field samples were collected, and RNA extracted by CEB, HCR, SA, AA, TR, and VR. AK led the mNGS, with support from VA, HCR, TR, CEB, CMT, and JLD. SH, GK, and CEB analyzed the resulting data. SH and CEB co-wrote the original draft of the manuscript, which all authors edited and approved.

## SUPPLEMENTARY MATERIAL

**Table S1:**
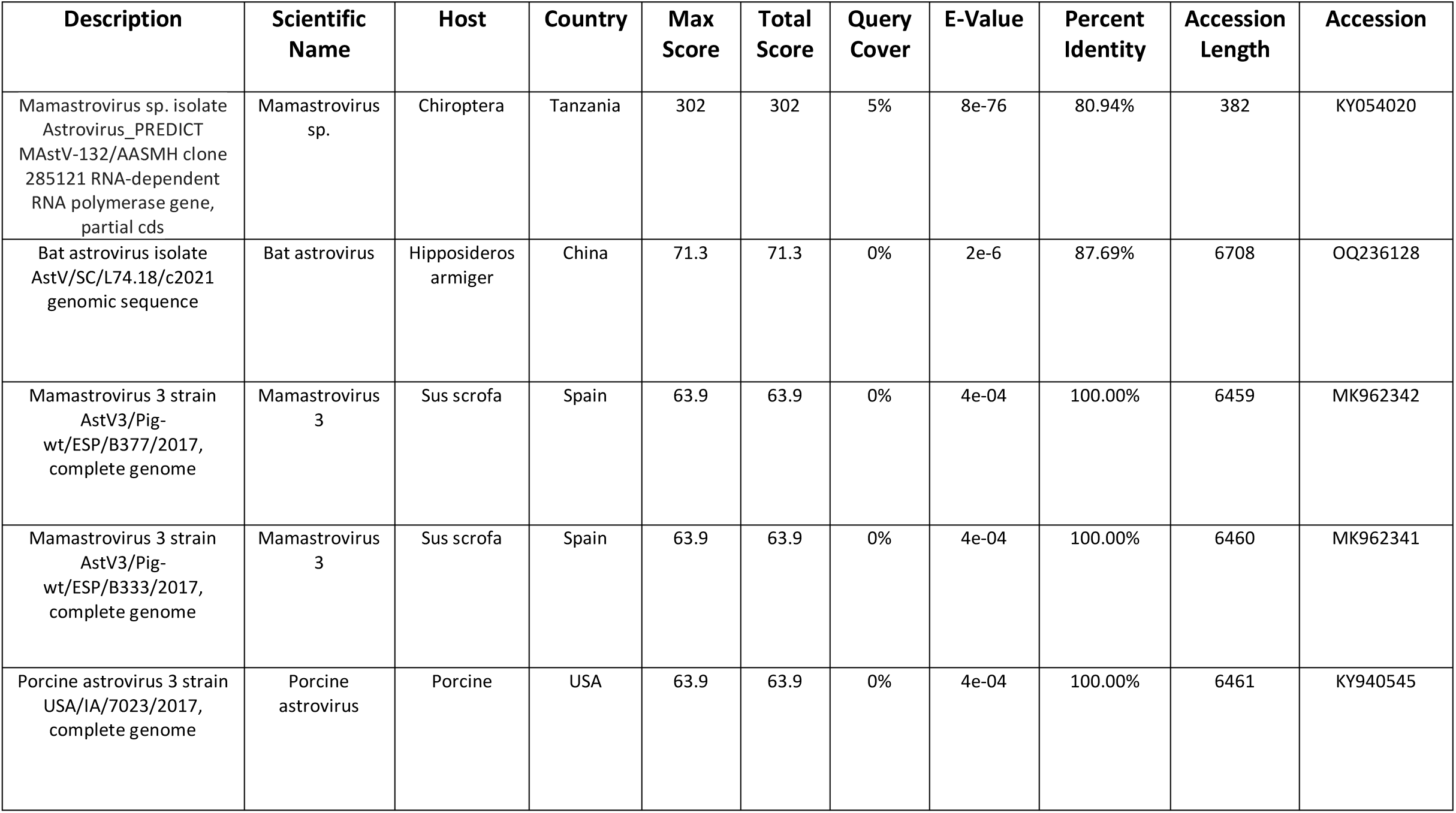
Top 5 NCBI BLASTn results (nucleotide-nucleotide) for novel near-full length astrovirus genome (accession number OQ606244).

| Description | Scientific Name | Host | Country | Max Score | Total Score | Query Cover | E-Value | Percent Identity | Accession Length | Accession |
| --- | --- | --- | --- | --- | --- | --- | --- | --- | --- | --- |
| Mamastrovirus sp. isolate<br>Astrovirus_PREDICT<br>MAstV-132/AASMH clone<br>285121 RNA-dependent<br>RNA polymerase gene,<br>partial cds | Mamastrovirus<br>sp. | Chiroptera | Tanzania | 302 | 302 | 5% | 8e-76 | 80.94% | 382 | KY054020 |
| Bat astrovirus isolate<br>AstV/SC/L74.18/c2021<br>genomic sequence | Bat astrovirus | Hipposideros<br>armiger | China | 71.3 | 71.3 | 0% | 2e-6 | 87.69% | 6708 | OQ236128 |
| Mamastrovirus 3 strain<br>AstV3/Pig-<br>wt/ESP/B377/2017,<br>complete genome | Mamastrovirus<br>3 | Sus scrofa | Spain | 63.9 | 63.9 | 0% | 4e-04 | 100.00% | 6459 | MK962342 |
| Mamastrovirus 3 strain<br>AstV3/Pig-<br>wt/ESP/B333/2017,<br>complete genome | Mamastrovirus<br>3 | Sus scrofa | Spain | 63.9 | 63.9 | 0% | 4e-04 | 100.00% | 6460 | MK962341 |
| Porcine astrovirus 3 strain<br>USA/IA/7023/2017,<br>complete genome | Porcine<br>astrovirus | Porcine | USA | 63.9 | 63.9 | 0% | 4e-04 | 100.00% | 6461 | KY940545 |

**Table S2:** Top 5 NCBI BLASTx results (nucleotide-amino acid) for novel near-full length astrovirus genome (accession number OQ606244).

| Description | Scientific Name | Host | Country | Max Score | Total Score | Query Cover | E-Value | Percent Identity | Accession Length | Accession |
| --- | --- | --- | --- | --- | --- | --- | --- | --- | --- | --- |
| ORF1a [Bat astrovirus] | Bat astrovirus | Eidolon helvum | Cameroon | 691 | 691 | 40% | 0.0 | 43.09% | 891 | AWV67084 |
| Nonstructural protein [Raccoon dog astrovirus 3] | Raccoon dog astrovirus 3 | Raccoon dog | China | 538 | 973 | 62% | 0.0 | 52.43% | 1362 | ULF48000 |
| ORF1ab [Porcine astrovirus 3] | Porcine astrovirus 3 | Sus scrofa | Japan | 518 | 939 | 61% | 0.0 | 51.88% | 1353 | BAX00201 |
| ORF1ab [Porcine astrovirus 3] | Porcine astrovirus 3 | Sus scrofa | Japan | 517 | 938 | 61% | 0.0 | 51.88% | 1353 | BAX00204 |
| ORF1ab [Porcine astrovirus 3] | Porcine astrovirus 3 | Sus scrofa | Japan | 514 | 936 | 61% | 0.0 | 52.49% | 1353 | BAX00207 |

**Table S3:** Presence/absence of astroviruses detected in Madagascar bats to date.

| Sub-Order | Family | Species | Detection | Citation |
| --- | --- | --- | --- | --- |
| Yinpteropchiroptera | Pteropodidae | *Pteropus rufus* | Yes | This paper |
|  |  | *Eidolon dupreanum* | Yes | This paper |
|  |  | *Rousettus madagascariensis* | Yes | This paper, Lebarbchenon et al |
|  | Rhinonycteridae | *Paratriaenops furculus* | Yes | Lebarbchenon et al, Hoarau et al |
|  |  | *Triaenops menamena* | Yes | Lebarbchenon et al, Hoarau et al |
|  | Hipposideridae | *Macronycteris commersoni* | Yes | Hoarau et al |
| Yangochiroptera | Molossidae | *Mormopterus jugularis* | No | Lebarbchenon et al, Hoarau et al |
|  |  | *Otomops madagascariensis* | No | Lebarbchenon et al, Hoarau et al |
|  |  | *Chaerephon atsinanana* | No | Hoarau et al |
|  |  | *Chaerephon leucogaster* | Yes | Hoarau et al |
|  |  | *Mops leucostigma* | Yes | Hoarau et al |
|  | Vespertilionidae | *Myotis goudoti* | Yes | Lebarbchenon et al, Hoarau et al |
|  |  | *Pipistrellus hesperidus* | No | Lebarbchenon et al, Hoarau et al |
|  | Miniopteridae | *Miniopterus gleni* | Yes | Lebarbchenon et al, Hoarau et al |
|  |  | *Miniopterus griveaudi* | Yes | Lebarbchenon et al, Hoarau et al |
|  |  | *Miniopterus mahafaliensis* | Yes | Hoarau et al |
|  |  | *Miniopterus soroculus* | Yes | Hoarau et al |
|  |  | *Miniopterus manavi* | Yes | Hoarau et al |

